# Connectome-wide structure-function coupling models implicate polysynaptic alterations in autism

**DOI:** 10.1101/2023.05.08.539817

**Authors:** Bo-yong Park, Oualid Benkarim, Clara F. Weber, Valeria Kebets, Serena Fett, Seulki Yoo, Adriana Di Martino, Michael P. Milham, Bratislav Misic, Seok-Jun Hong, Boris C. Bernhardt

## Abstract

Autism spectrum disorder (ASD) is one of the most common neurodevelopmental diagnoses. Although incompletely understood, structural and functional network alterations are increasingly recognized to be at the core of the condition. We utilized multimodal imaging and connectivity modeling to study structure-function coupling in ASD, and probed mono- and polysynaptic mechanisms on structurally-governed network function. We examined multimodal magnetic resonance imaging data in 47 ASD and 37 neurotypical controls from the Autism Brain Imaging Data Exchange (ABIDE) II initiative. We predicted intrinsic functional connectivity from structural connectivity data in each participant using a Riemannian optimization procedure that varies the times that simulated signals can unfold along tractography-derived personalized connectomes. In both ASD and neurotypical controls, we observed improved structure-function prediction at longer diffusion time scales, indicating better modeling of brain function when polysynaptic mechanisms are accounted for. Prediction improvements were marked in transmodal association systems, such as the default mode network, in both controls and ASD. Improvements were, however, lower in ASD in a polysynaptic regime at higher simulated diffusion times. Regional differences followed a sensory-to-transmodal cortical hierarchy, with an increased gap between groups in transmodal compared to sensory/motor systems. Multivariate associative techniques revealed that structure-function differences reflected inter-individual differences in autistic symptoms and verbal as well as non-verbal intelligence. Our network modeling approach sheds light on atypical structure-function coupling in autism, and suggests that polysynaptic network mechanisms are implicated in the condition and can help explain its wide range of associated symptoms.

## Introduction

Autism spectrum disorder (ASD) is a common neurodevelopmental diagnosis encompassing atypical social and communication abilities, repetitive behaviors and interests, and sometimes altered sensory and perceptual processing as well as imbalances in verbal and non-verbal abilities (Christensen et al., 2018; Mottron et al., 2006). While biological underpinnings remain incompletely understood, convergent evidence supports a key role of atypical brain networks. Indeed, there is now an increasing catalog of ASD-related genes and pathways involved in synaptic and circuit organization (Geschwind, 2011; Quesnel-Vallières et al., 2019; Rylaarsdam and Guemez-Gamboa, 2019). Moreover, several histopathological studies suggest dendritic reorganization (Hutsler and Zhang, 2010; Martínez-Cerdeño, 2017), alterations in cortical lamination (Hutsler et al., 2007; Simms et al., 2009), and atypical columnar layout in individuals with ASD (Amaral et al., 2008; McKavanagh et al., 2015). Molecular and circuit findings are complemented by *in vivo* magnetic resonance imaging (MRI) studies, suggesting atypical structural and functional network organization, often pointing to a mosaic pattern of increased and decreased connectivity in ASD. Recent studies have represented structural and functional network organization in compact connectivity spaces, identified via unsupervised dimensionality reduction techniques, and tracked typical and atypical development (S.-J. Hong et al., 2019; Huntenburg et al., 2018; Margulies et al., 2016; Park et al., 2022, 2021b; Tian et al., 2020). In neurotypical adults, these techniques have robustly identified main spatial axes corresponding to the functional cortical hierarchy, differentiating sensory and motor systems interacting with the outside world from transmodal networks, such as default-mode and limbic networks, implicated in higher-order and social cognition (Margulies et al., 2016). Translating this framework to ASD, increasing evidence suggests a reduced hierarchical differentiation between sensory/motor and transmodal systems both at the level of structural and functional connectivity, which have been shown to relate to autism risk gene expression patterns (Park et al., 2021b). Overall, findings suggest that ASD perturbs neural circuit organization across multiple, likely interacting spatial scales.

A key assumption of neuroscience is that brain structure and function are intertwined. Expanding from experimental explorations in non-human animals, imaging studies in neurotypical populations have addressed structure-function coupling in the living human brain (Baum et al., 2020; Honey et al., 2009; Miŝic et al., 2016; Park et al., 2021d; Snyder and Bauer, 2019; Suárez et al., 2020; Vázquez-Rodríguez et al., 2019). Generally, such work seeks to identify a mapping from structural connectivity (approximated via diffusion MRI tractography) to functional connectivity (estimated via functional MRI signal correlations). Approaches include statistical associative techniques, biophysical modeling, and graph communication models (Avena-Koenigsberger et al., 2019, 2018; Bazinet et al., 2021; Becker et al., 2018; Breakspear, 2017; Deco et al., 2013; Goñi et al., 2014; Honey et al., 2009; Miŝic et al., 2016; Rosenthal et al., 2018; Seguin et al., 2018; Wang et al., 2019). This body of work emphasizes that functional interactions unfold both along direct monosynaptic connections as well as indirect polysynaptic pathways (Damoiseaux and Greicius, 2009; Goñi et al., 2014; Honey et al., 2009; Seguin et al., 2019; Suárez et al., 2020). In neurotypical adults, our team recently proposed a novel approach to simulate functional interactions from structural connectivity with high fidelity and at an individual-participant level (Benkarim et al., 2022). This work derived low-dimensional eigenspaces from a structural connectome, on which virtual signal diffusion models were then used to predict inter-regional functional interactions. These diffusion processes unfold along existing connections and are governed by a free diffusion time parameter, with higher diffusion times implicating an increasing contribution of indirect pathways to functional interactions. In other words, this Riemannian manifold optimization framework can parameterize the impact of polysynaptic communication on global structure-function coupling. At a regional scale, comparing simulations with empirically measured data showed that while functional interactions of sensory and motor systems can be adequately modeled with only a limited number of synaptic steps, accurate simulations of interactions of transmodal systems require longer time scales, and thus a more polysynaptic regime. As such, mono- and polysynaptic communication mechanisms underpinning structure-function coupling in healthy individuals can be compactly described along a unimodal to transmodal brain hierarchy.

Our study examined structure-function relations in autism and explored the differential impact of mono- *vs* polysynaptic communication. Core to our approach was a Riemannian optimization and modeling framework (Benkarim et al., 2022), which has shown state-of-the-art performance in predicting functional interactions from structural connectivity data in single neurotypical individuals. We studied global and region-specific differences in prediction accuracy across diffusion times in individuals with ASD and neurotypicals to evaluate the impact of mono- and polysynaptic network communication. The topography of ASD-related alterations was spatially associated with canonical features of macroscale functional organization, namely intrinsic functional systems and sensory-transmodal cortical hierarchical gradients. Using partial least squares regression, we finally associated ASD-related alterations with autistic symptoms and measures of verbal/non-verbal intelligence to explore how atypical structure-function coupling reflects behavioral phenotypes.

## Results

### Global imbalances in structure-function coupling in ASD

Based on connectome manifold models (Benkarim et al., 2022), we simulated resting-state functional connectivity among 200 cortical regions (Schaefer et al., 2018) from tractography-derived structural connectivity data (Benkarim et al., 2022). In brief, the technique *(i)* applies nonlinear dimensionality reduction (*i.e.,* diffusion map embedding) (Coifman and Lafon, 2006; Vos de Wael et al., 2020) to a structural connectome, *(ii)* varies the diffusion time parameter *t* of the embedding technique to simulate connectivity-guided random walks (**Fig. 1A**), and *(iii)* the kernels derived from the corresponding diffusion times using a radial basis function are fused to minimize the difference between the actual functional connectivity and diffusion maps applied. Before generating the kernels, the algorithm uses a transformation matrix to rotate diffusion maps to find optimal paths through which to propagate information between different brain regions at each diffusion time. Structure-function coupling was quantified as the linear correlation between empirical and simulated functional connectivity matrices across diffusion times *t* (from *t* = 1 to *t* = 10, with higher *t* indicating an increased contribution of polysynaptic communication across indirect paths; **Fig. 1B**). In both neurotypicals and ASD, coupling monotonically increased with higher diffusion times. Notably, controls showed higher prediction performance between *t* = 2–4. We quantitatively assessed between-group differences in coupling using 1,000 permutation tests by shuffling subject indices, and confirmed higher performance in controls relative to ASD between *t* = 2 and *t* = 4 (p_perm_ = 0.020, 0.044, 0.038; **Fig. 1C**). The results indicate that both controls and ASD displayed an influence of polysynaptic communication on structure-function coupling, and stronger global coupling in controls than in ASD.

**Fig. 1.**
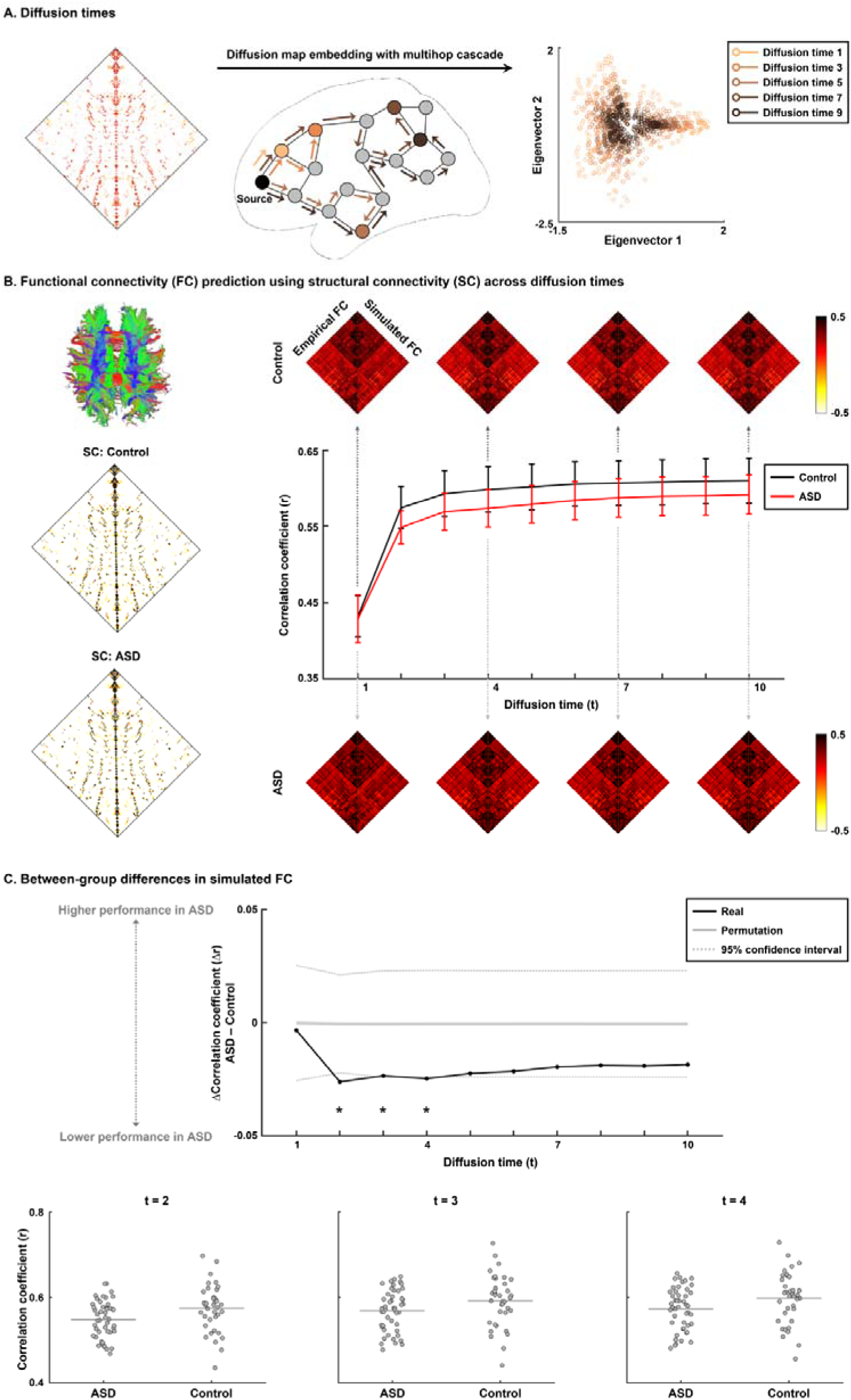
Global imbalances in structure-function coupling in ASD. **(A)** Schema of the Riemannian manifold optimization approach that was used to simulate functional connectivity (FC) along a structural connectome (SC) as a function of diffusion time *t*. **(B)** Group-level SC matrices in controls and ASD (*left*). Correlation coefficients between empirical and simulated FC in controls (*dark gray*) and ASD (*light gray*) as a function of *t* (*right*). Error bars represent the SD across individuals. Shown are empirical (*left*) and simulated FC (*right*) matrices across four representative diffusion times (*t* = 1, 4, 7, and 10). **(C)** Between-group differences in prediction performance between controls and ASD (*upper panel*). A black line indicates real differences in prediction performance between groups, a solid gray line indicates mean prediction accuracy differences across 1,000 permutation tests, and dotted gray lines indicate the 95% confidence interval. Significant between-group differences are reported with asterisks. Shown are correlation coefficients between an individual’s empirical and simulated FC for those diffusion times that showed significant between-group differences (*lower panels*). *Abbreviations*: ASD, autism spectrum disorder; SD, standard deviation.

### Regional structure-function imbalances

We assessed regional prediction performance gains across variable diffusion times *t* to explore the contribution of polysynaptic communication on the prediction of brain function. For both controls and ASD, sensory/motor areas showed higher prediction accuracy at low diffusion times compared to transmodal systems (*i.e.,* default-mode network and paralimbic cortices). With increasing diffusion times, global prediction performance increased in both groups, with higher performance in controls (**Fig. 2A**). To assess improvements in prediction accuracy across diffusion times, we calculated prediction accuracy differences between *t* = 10 and *t* = 1 (Δprediction accuracy) in both cohorts separately (**Fig. 2B**). We observed marked improvements (false discovery rate (FDR) < 0.05) in transmodal compared to sensory/motor systems, and improvements were larger in controls than ASD.

**Fig. 2.**
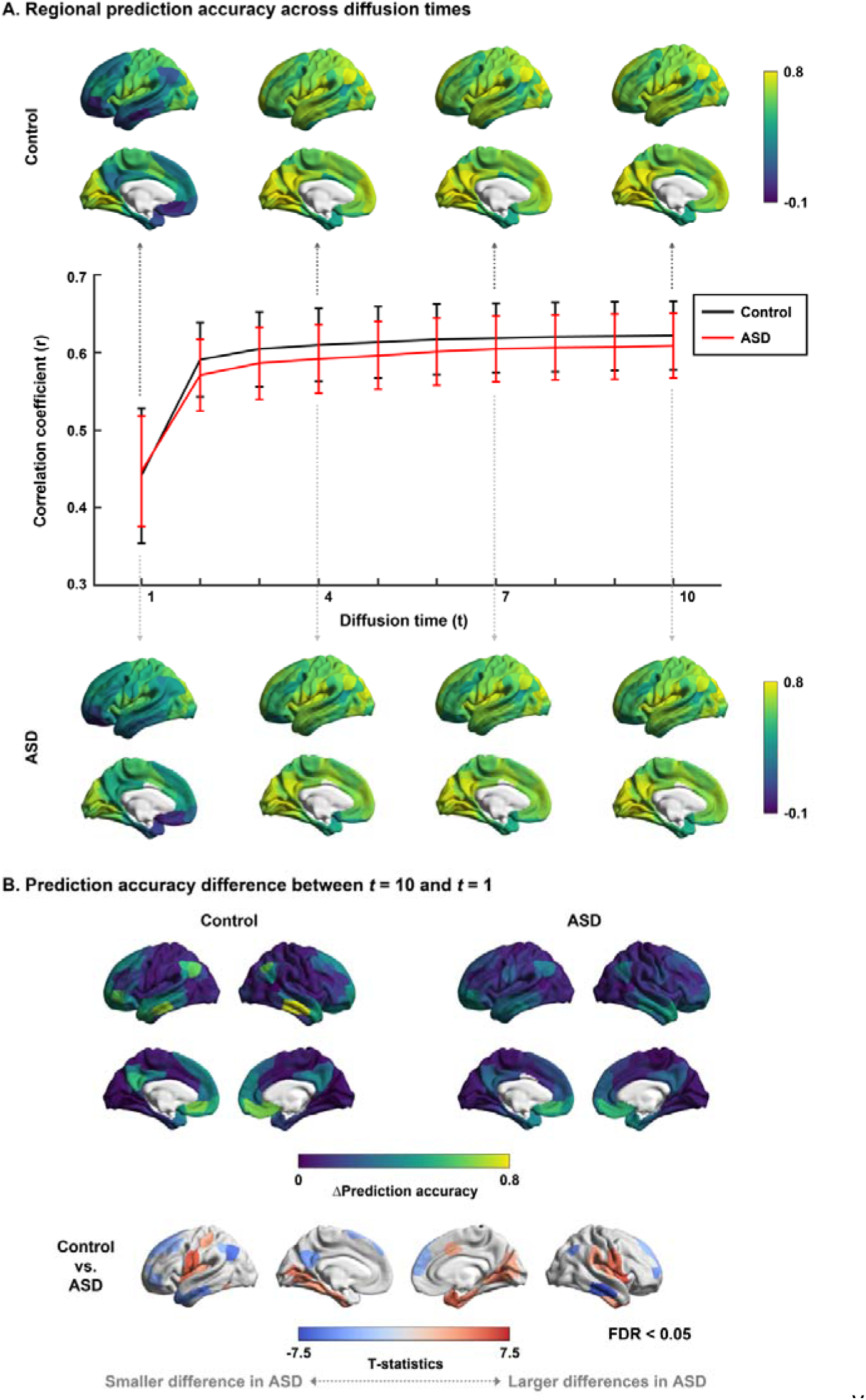
Regional structure-function imbalances. **(A)** Correlation coefficients between empirical and simulated FC across different diffusion times *t* are shown on brain surfaces for control and ASD groups. The plot indicates correlation coefficients between the empirical and simulated FC in controls (*dark gray*) and ASD (*light gray*) as a function of diffusion time. Error bars represent the SD across brain regions. **(B)** Shown are differences in prediction accuracy between the highest (*t* = 10) and lowest (*t* = 1) diffusion times (Δprediction accuracy) for both groups (*upper panels*). We assessed between-group differences in Δprediction accuracy between controls and ASDs (*lower panel*). *Abbreviations*: ASD, autism spectrum disorder; SD, standard deviation; FDR, false discovery rate.

### Topographic associations to structure-function imbalances

We stratified findings with respect to established taxonomies of intrinsic functional organization. First, we related our findings to a prior atlas that decomposes the cortex into seven intrinsic systems (Yeo et al., 2011) and to a foundational taxonomy that subdivides the cortex into four hierarchical levels (Mesulam, 1998) (**Fig. 3A**). In particular, we assessed across-diffusion time improvements in structure-function prediction performance (Δprediction accuracy) as a function of the intrinsic system and or hierarchical level, and noted smaller improvement in default-mode and frontoparietal networks in ASD relative to controls, while sensory and attention networks showed increased improvement. Second, we explored associations with the principal functional gradient, which discriminates sensory/visual from transmodal systems in a continuous manner based on data-driven connectome analysis. The first principal functional gradient was estimated from resting-state functional connectivity obtained from the Human Connectome Project database (Van Essen et al., 2013), using the BrainSpace toolbox version 0.1.10 (https://github.com/MICA-MNI/BrainSpace) (Coifman and Lafon, 2006; Vos de Wael et al., 2020) (**Fig. 3B**). We observed significant correlations with the functional gradient, even after accounting for spatial autocorrelation, in both groups (control: r = 0.597 ± 0.103, p_spin_ < 0.001; ASD: r = 0.441 ± 0.200, p_spin_ = 0.021; **Fig. 3B**). Associations were significantly different between groups, and stronger in controls (two-sample t-tests with 1,000 permutations p < 0.001; **Fig. 3B**).

**Fig. 3.**
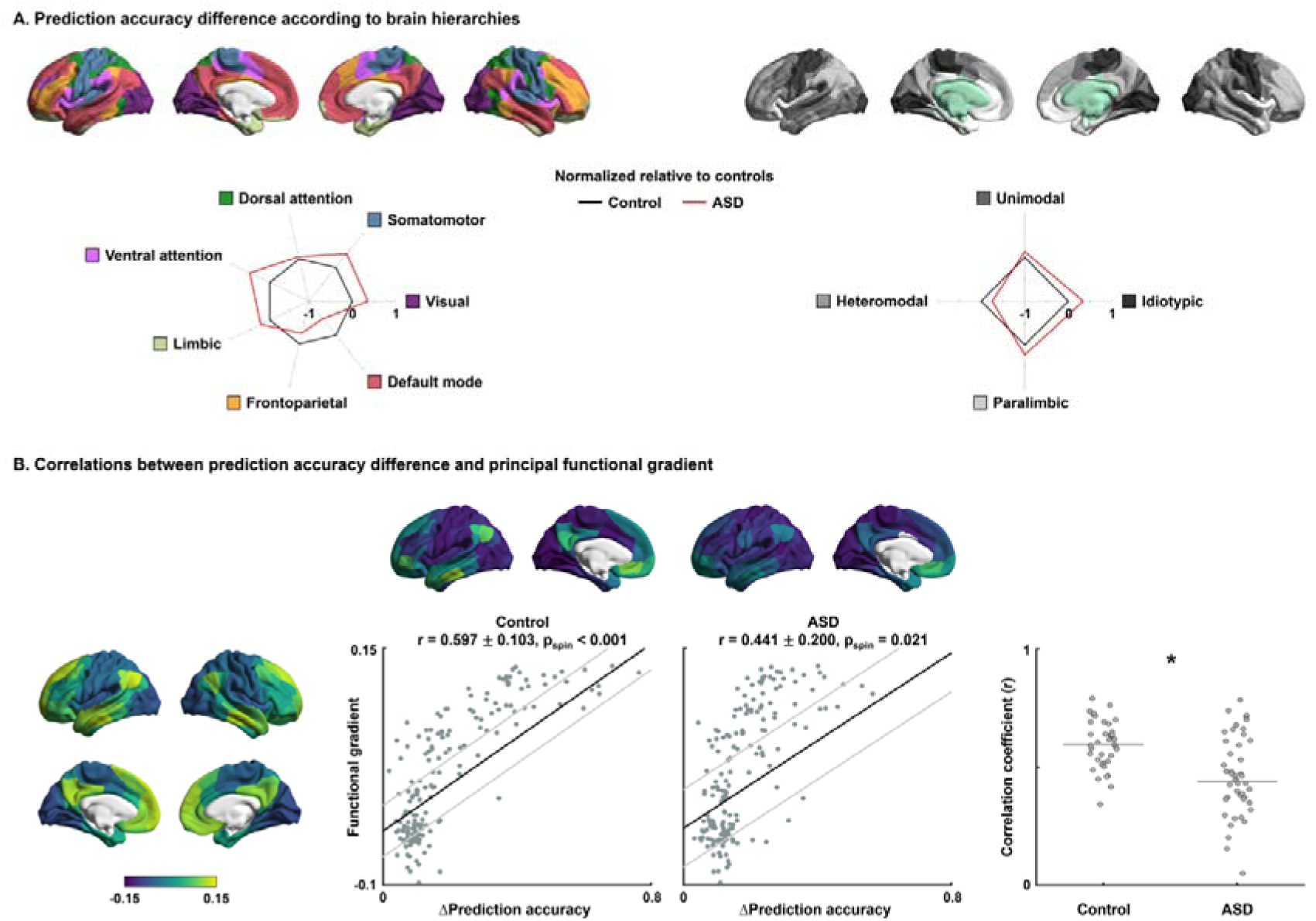
Topographic associations. **(A)** We stratified the prediction accuracy difference between diffusion time *t = 10* and *t = 1* (Δprediction accuracy) according to functional communities (*left*) (Yeo et al., 2011) and cortical hierarchies (*right*) (Mesulam, 1998). Spider plots show normalized Δprediction accuracy, where the values of ASD are normalized relative to controls. **(B)** The principal functional gradient is visualized on brain surfaces (*left*). We calculated linear correlations between the gradient and Δprediction accuracy for both controls and ASD individuals, where the gray lines indicate SD across individuals (*center*). In the right panel, one can see the correlation coefficients of each individual in both groups, where the asterisk indicates a significant difference. *Abbreviations*: ASD, autism spectrum disorder; SD, standard deviation.

### Associations with behavioral phenotypes

We studied associations between prediction accuracy difference across diffusion times (Δprediction accuracy) and behavioral phenotypes of ADOS scores (social cognition, communication, and repetitive behavior) as well as verbal and non-verbal intelligence quotient (IQ) and their ratio (verbal/non-verbal IQ) (Hong et al., 2022) using partial least squares (PLS) analyses (Krishnan et al., 2011; McIntosh and Mišić, 2013) (see *Methods*). We performed the PLS analysis with 1,000 bootstraps, and the first latent variable explained 33.4 % of covariance between Δprediction accuracy and behavioral phenotypes (**Fig. 4A**). The estimated PLS scores showed significant correlations across bootstraps (r = 0.426 ± 0.093, p_perm_ = 0.010; **Fig. 4B**). We assessed the contribution of these features using bootstrap ratio calculated based on the loadings (Zeighami et al., 2019) (see *Methods*). We found that improvement of prediction accuracy in sensory and frontoparietal regions was associated with lower social cognition and communication-related autistic symptoms and IQ ratio, indicating less autistic characteristics (**Fig. 4C**). On the other hand, prediction accuracy improvement in temporal and limbic regions was associated with higher autistic symptom, particularly, repetitive behaviors (**Supplementary Fig. 1**).

**Fig. 4.**
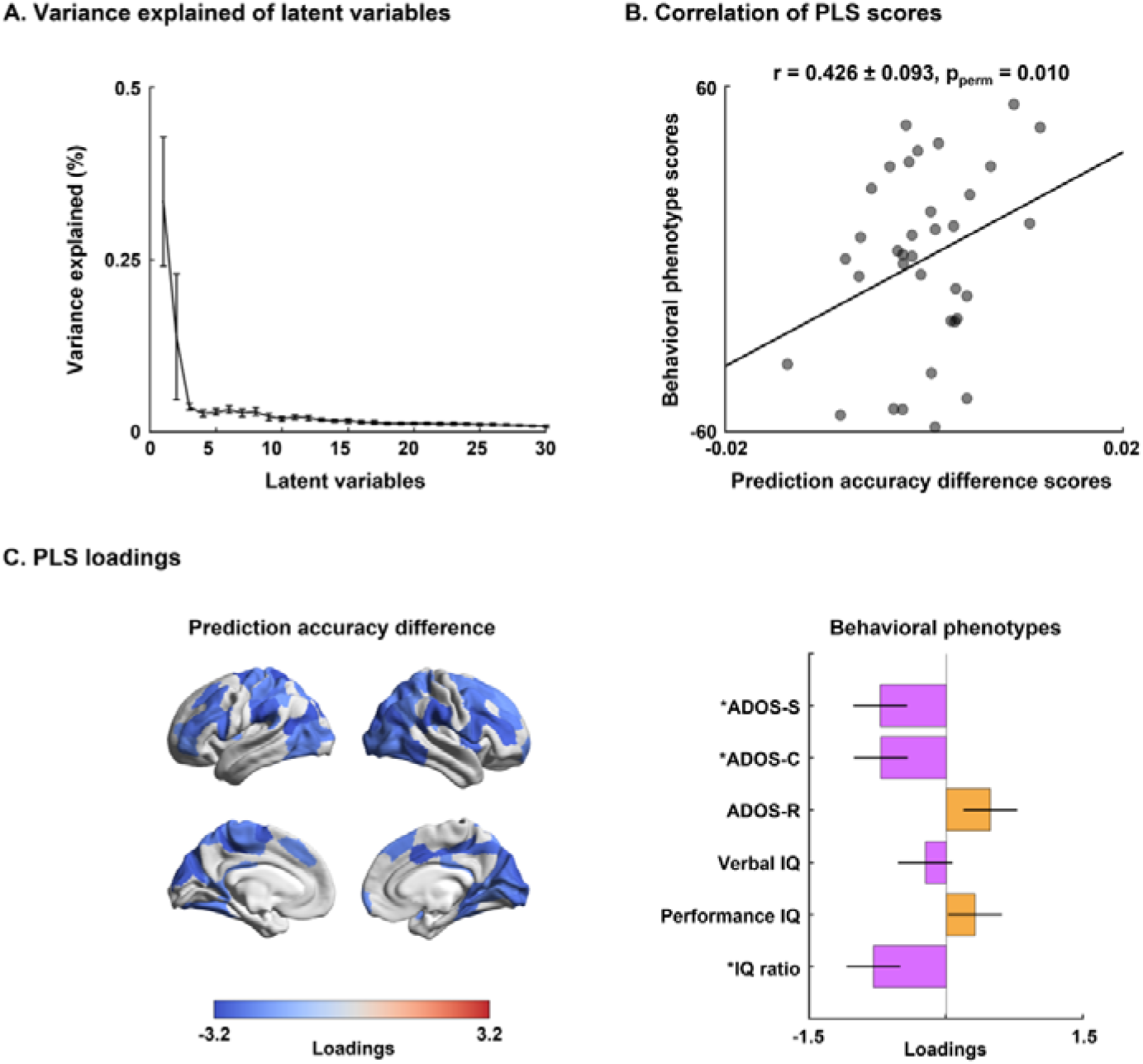
Multivariate associations between structure-function coupling imbalances and behavioral phenotypes of ASD. **(A)** The scree plot shows the percent variance explained by each latent variable, where the error bars indicate SD across bootstraps. **(B)** We calculated linear correlations between PLS scores of Δprediction accuracy and behavioral phenotypes of the first latent variable in ASD, which explained almost 33.4% of the variance. **(C)** Shown are PLS loading-based bootstrap ratios of Δprediction accuracy (*left*) and behavioral phenotypes (*right*). Brain regions and behavioral phenotypes that showed significance are shown and marked with asterisks. *Abbreviations*: SD, standard deviation; PLS, partial least squares; ADOS-S, Autism Diagnostic Observation Schedule – social cognition; ADOS-C, Autism Diagnostic Observation Schedule – communication; ADOS-R, Autism Diagnostic Observation Schedule – repeated behavior; IQ, intelligence quotient.

### Sensitivity analyses

*a) Parcellation schemes.* In addition to the 200 cortical regions, we found consistent results when dividing the cortex into 100 and 300 regions (**Supplementary Figs. 2-3**).
*b) Only male participants.* We additionally performed the same analyses using only male participants and found comparable results (**Supplementary Fig. 4**).
*c) Site effects.* We performed the analyses for each site separately. We found overall similar patterns but decreased effects, which may be due to the small sample size (**Supplementary Fig. 5**).

## Discussion

The correspondence of brain structure and function is a tenet of neuroscience (Baum et al., 2020; Honey et al., 2009; Miŝic et al., 2016; Park et al., 2021d; Snyder and Bauer, 2019; Suárez et al., 2020; Vázquez-Rodríguez et al., 2019), and the advent of multimodal imaging and connectomics methods has culminated in multiple efforts to predict large-scale brain function and inter-regional functional interactions from descriptions of brain wiring in the healthy human brain (Benkarim et al., 2022; Damoiseaux and Greicius, 2009; Goñi et al., 2014; Honey et al., 2009; Seguin et al., 2019; Suárez et al., 2020). Here, we utilized unsupervised connectivity manifold learning and alignment techniques to index structure-function coupling in ASD and to explore the role of polysynaptic communication mechanisms. Studying individuals with ASD and neurotypical controls, we observed structure-function coupling in both groups to be overall high and generally increasing when additionally incorporating polysynaptic communication, particularly in transmodal systems. On the other hand, ASD showed reduced structure-function coupling compared to controls, in particular in polysynaptic regimes and transmodal regions. Structure-function coupling imbalances in ASD were aligned with prototypical descriptions of the primate cortical hierarchy, indicating a sensory-to-transmodal gradient of alterations in structure-function coupling in ASD. Findings reflected autism symptoms and imbalances in verbal/non-verbal intelligence dimensions. Collectively, our findings show hierarchy-dependent imbalances in structurally-governed network communication in ASD, and may offer a novel and behaviorally relevant perspective of autism connectopathy.

Our work investigated connectome-level structure-function coupling using a Riemannian manifold optimization procedure (Benkarim et al., 2022). In a recent study in neurotypical adults, this approach provided a faithful individual participant-level prediction of intrinsic functional interactions based on structural connectomes. It can be tuned across diffusion time parameters, interpretable as an increasing influence of polysynaptic structure-function coupling mechanisms. Comparing prediction accuracy between neurotypicals and ASD, our findings revealed globally reduced coupling in the latter. Coupling was particularly reduced towards higher diffusion times, and ASD-related reductions were most marked in transmodal systems such as the default mode and frontoparietal networks. Such findings indicate a hierarchy-dependent alteration in structure-function coupling in ASD, particularly in polysynaptic subnetworks. These findings suggest that links between brain structure and function are not as straightforward in ASD compared to controls, which may relate to several previously identified factors. Neuroimaging studies have shown atypical cortical morphology and microstructure, aberrant white matter fiber architecture, and reorganized structural network topology in ASD (Cai et al., 2022; Hong et al., 2018; S. J. Hong et al., 2019). Despite only a little work assessing links between structural alterations and atypical function in ASD, studies have indicated atypical functional connectivity between different brain areas (Di Martino et al., 2014; Hull et al., 2017; Müller et al., 2011). Moreover, several reports emphasized increased spatial shifting of functional network layout in ASD, a finding also referred to as idiosyncrasy (Benkarim et al., 2021), alongside findings suggesting increased signal variation in this cohort (Takahashi et al., 2016). These factors may collectively result in lower predictability of functional signaling and interactions from structural connectivity information, and hence contribute to the observed findings in this study.

Brain hierarchy along the sensory/motor-association axis underpins primate cortical organization, initially inferred from invasive *post-mortem* findings in non-human animals (Mesulam, 1998). Recently, our understanding of the cortical hierarchical organization has been solidified with human neuroimaging, notably functional connectivity research (Bethlehem et al., 2020; Margulies et al., 2016; Mckeown et al., 2020; Murphy et al., 2019; Park et al., 2021c), microstructural profiling (Burt et al., 2018; Paquola et al., 2019), and tractography-derived structural connectomics (Kharabian Masouleh et al., 2020; Park et al., 2021a, 2021b). In our study, inter-regional variations in structure-function prediction performance followed dimensional and clustering-based approximations of the cortical functional hierarchy. In particular, we observed lower coupling towards transmodal systems when incorporating monosynaptic mechanisms, which, however, increased with larger diffusion times and hence polysynaptic communication. Overall reduced structure-function coupling in transmodal systems compared to sensory/motor and unimodal networks echoes prior findings (Valk et al., 2022; Vázquez-Rodríguez et al., 2019), in particular when bare diffusion MRI tractography measures without explicit incorporation of polysynaptic communication inform the modeling strategy. Transmodal regions are known to increasingly engage in long-range and more centralized communication, underpinning integrative cognitive functions (Park et al., 2021d). In our work, performance reductions in ASD relative to neurotypicals related mainly to reduced hierarchy-specific gains in predicting functions that would have otherwise resulted from the incorporation of polysynaptic communication in ASD. Previous work from our group and others based on functional and structural neuroimaging has suggested atypical connectome hierarchy, and suggested that densely integrated rich core nodes may assume a major role in this process (S.-J. Hong et al., 2019; Park et al., 2021b), possibly in lieu of their implication in multiple, polysynaptic communication pathways.

Multivariate associative techniques revealed that altered structure-function relations in ASD reflected behavioral symptoms and cognitive phenotypes, here indexed by the ADOS scale and verbal and non-verbal intelligence dimensions (Di Martino et al., 2011; Hong et al., 2022; S.-J. Hong et al., 2019; Park et al., 2021b). It should be noted that our results were derived from small samples and assessed using four-fold cross-validation only, requiring validations in larger samples with multimodal imaging data to assess generalizability. Findings, nevertheless, suggested a broad implication of different brain systems, notably transmodal systems, such as the default-mode network. These systems have been shown to contribute to both typical and atypical social interaction and communication, and higher cognitive processes more generally (Assaf et al., 2010; Mars et al., 2012; Padmanabhan et al., 2017; Paquola et al., 2022; Raichle, 2015; Smallwood et al., 2021). Moreover, systems at the apex of the putative cortical hierarchy undergo ongoing maturational processes in typical childhood and adolescence, which shift networks towards a more clustered layout and progressively differentiate these from other macroscale networks, possibly due to the strengthening of long-range connections (Baum et al., 2020; Fan et al., 2021; Park et al., 2022). Our findings suggest that atypical polysynaptic communication in higher-order transmodal areas, in part, reflects those symptoms, and could serve as a potential diagnostic marker of affected individuals.

## Methods

### Study participants

We studied 84 participants (47 ASD, 37 neurotypicals) obtained from two independent sites of (1) New York University Langone Medical Center (NYU) and (2) Trinity College Dublin (TCD) from the Autism Brain Imaging Data Exchange initiative (ABIDE-II; https://fcon_1000.projects.nitrc.org/indi/abide) (Di Martino et al., 2017). Inclusion criteria were: (i) sites included children and adults with autism and controls with ≥10 individuals per group, (ii) multimodal MRI data (*i.e.,* T1-weighted, rs-fMRI, and diffusion MRI) available, (iii) acceptable cortical surface extraction on T1-weighted MRI, (iv) low head motion in the rs-fMRI time series (*i.e.,* >0.3 mm framewise displacement). Individuals with ASD were diagnosed by an in-person interview with clinical experts and gold standard instruments from the Autism Diagnostic Observation Schedule (ADOS) (Lord et al., 2000) and/or Autism Diagnostic Interview-Revised (ADI-R) (Lord et al., 1994). Neurotypical controls did not have any history of mental disorders. For all groups, participants who had genetic disorders associated with autism (*i.e.,* Fragile X), contraindications to MRI scanning, and who were pregnant were excluded. Detailed demographic information of the participants is reported in **Supplementary Table 1**. ABIDE data collections were performed in accordance with local Institutional Review Board guidelines. In accordance with HIPAA guidelines and 1000 Functional Connectomes Project/INDI protocols, all ABIDE datasets have been fully anonymized, with no protected health information included.

### MRI acquisition

We obtained the data from two independent sites.

i. NYU: Imaging data were acquired using a 3T Siemens Allegra scanner. The T1-weighted data were obtained using a 3D magnetization prepared rapid acquisition gradient echo (MPRAGE) sequence (repetition time (TR) = 2,530 ms; echo time (TE) = 3.25 ms; inversion time (TI) = 1,100 ms; flip angle = 7°; matrix = 256 × 192; and voxel size = 1.3 × 1.0 × 1.3 mm^3^). The rs-fMRI data were acquired using a 2D echo planar imaging (EPI) sequence (TR = 2,000 ms; TE = 15 ms; flip angle = 90°; matrix = 80 × 80; number of volumes = 180; and voxel size = 3.0 × 3.0 × 4.0 mm^3^). The diffusion MRI data were obtained using a 2D spin echo EPI (SE-EPI) sequence (TR = 5,200 ms; TE = 78 ms; matrix = 64 × 64; voxel size = 3 mm^3^ isotropic; 64 directions; b-value = 1,000 s/mm^2^; and 1 b0 image).
ii. TCD: Imaging data were acquired using a 3T Philips Achieva scanner. The T1-weighted MRI were acquired using a 3D MPRAGE (TR = 8,400 ms; TE = 3.90 ms; TI = 1,150 ms; flip angle = 8°; matrix = 256 × 256; voxel size = 0.9 mm^3^ isotropic). The rs-fMRI data were acquired using a 2D EPI (TR = 2,000 ms; TE = 27 ms; flip angle = 90°; matrix = 80 × 80; number of volumes = 210; and voxel size = 3.0 × 3.0 × 3.2 mm^3^). The diffusion MRI data were acquired using a 2D SE-EPI (TR = 20,244 ms; TE = 79 ms; matrix = 124 × 124; voxel size = 1.94 × 1.94 × 2 mm^3^; 61 directions; b-value = 1,500 s/mm^2^; and 1 number b0 image).

### Data preprocessing

We preprocessed the T1-weighted data using FreeSurfer version 6.0 (Dale et al., 1999; Fischl, 2012; Fischl et al., 2001, 1999a, 1999b; Ségonne et al., 2007), which includes gradient nonuniformity correction, skull stripping, intensity normalization, and tissue segmentation. White and pial surfaces were generated through triangular surface tessellation, topology correction, inflation, and spherical registration to the fsaverage template surface. The rs-fMRI data were previously processed using C-PAC (https://fcp-indi.github.io) (Craddock et al., 2013), and provided by the ABIDE database (http://preprocessed-connectomes-project.org/abide/). The pipeline included slice timing and head motion correction, skull stripping, and intensity normalization. Nuisance variables of head motion, average white matter and cerebrospinal fluid signal, and linear/quadratic trends were removed using CompCor (Behzadi et al., 2007). Band-pass filtering between 0.01 and 0.1 Hz was applied, and rs-fMRI data were co-registered to T1-weighted data in MNI152 standard space with boundary-based rigid-body and nonlinear transformations. The rs-fMRI data were mapped to subject-specific midthickness surfaces and resampled to Conte69. Finally, surface-based spatial smoothing with a full-width-at-half-maximum of 5 mm was applied. The diffusion MRI data were processed using Mrtrix3 (Tournier et al., 2019), including correction for susceptibility distortions, head motion, and eddy currents. Quality control involved visual inspection of T1-weighted data, and cases with faulty cortical segmentation were excluded. Cases with an rs-fMRI data framewise displacement >0.3 mm were also excluded (Power et al., 2014, 2012).

### Structural and functional connectivity

Structural connectomes were generated from preprocessed diffusion MRI data using Mrtrix3 (Tournier et al., 2019). Anatomical constrained tractography was performed using different tissue types derived from the T1-weighted image, including cortical and subcortical grey matter, white matter, and cerebrospinal fluid (Smith et al., 2012). The T1-weighted was registered to the diffusion MRI data with boundary-based registration, and the transformation was applied to different tissue types to register them onto the native diffusion MRI space. Multi-shell and multi-tissue response functions were estimated (Christiaens et al., 2015), and constrained spherical deconvolution and intensity normalization were performed (Jeurissen et al., 2014). Seeding from all white matter voxels, the tractogram was generated based on a probabilistic approach (Tournier et al., 2010, 2019, 2012) with 40 million streamlines, with a maximum tract length of 250 and a fractional anisotropy cutoff of 0.06. Subsequently, spherical-deconvolution informed filtering of tractograms (SIFT2) was applied to reconstruct whole-brain streamlines weighted by the cross-section multipliers, which considers the fiber bundle’s total intra-axonal space across its full cross-sectional extent (Smith et al., 2015). The structural connectome was built by mapping the reconstructed cross-section streamlines onto the Schaefer atlas with 200 parcels (Schaefer et al., 2018), then log-transformed to adjust for the scale (Fornito et al., 2016). Functional connectivity matrices were generated by calculating Pearson’s correlations of time series between two different brain regions defined using the Schaefer atlas with 200 parcels (Schaefer et al., 2018), and the correlation coefficients were Fisher’s r-to-z transformation to render data more normally distributed (Thompson and Fransson, 2016).

### Functional connectivity prediction using structural connectivity

To predict functional connectivity from structural connectivity, we opted for a recently introduced Riemannian optimization approach (Benkarim et al., 2022). Core to this approach is the application of diffusion map embedding, a nonlinear dimensionality technique, to the structural connectivity matrix to generate low-dimensional eigenvectors (*i.e.,* diffusion maps) (Coifman and Lafon, 2006). The diffusion maps are controlled by diffusion time *t*, which controls the scale of eigenvalues. If the diffusion time increases, the brain regions are located more closely on the low-dimensional eigenspace (**Fig. 1A**). We, thus, can approximate the distances of eigenvectors between different regions with varying diffusion times. To predict functional connectivity, this approach uses kernel fusion to find a weighted combination of the kernels derived from the diffusion maps at each diffusion time using a radial basis function. The approach further uses a transformation matrix to rotate the diffusion maps before computing the kernels for each diffusion time, where the rotation matrix may help identify the optimal paths through which to propagate information between different brain regions. Here, to solve the optimization problem, the Riemannian conjugate gradient algorithm (Absil et al., 2008) implemented in the Pymanopt toolbox (Townsend et al., 2016) was used. Details can be found elsewhere (Benkarim et al., 2022). We performed the prediction procedure for the ASD and control groups separately with a five-fold cross-validation. The prediction performance was assessed by calculating Pearson’s correlation of the upper triangular elements between empirical and predicted functional connectivity matrices (**Fig. 1B**). We assessed the prediction accuracy with varying diffusion times between *t* = 1 and *t* = 10. Differences in prediction accuracy between ASD and control groups were assessed based on 1,000 permutation tests (**Fig. 1C**). We randomly assigned subject indices and calculated differences in prediction accuracy between the new groups (Δr) to construct a null distribution. If the real difference did not belong to 95% of the null distribution, it was deemed significant.

### Between-group differences in regional prediction accuracy across diffusion times

We also assessed regional prediction performances across varying diffusion times. For each diffusion time, we calculated Pearson’s correlations between the empirical and predicted functional connectivity matrix of each brain region (**Fig. 2A**). To assess the improvement of the prediction accuracy across diffusion times, we calculated the difference in prediction accuracy between the highest (*t* = 10) and lowest (*t* = 1) diffusion times (i.e., Δprediction accuracy) (**Fig. 2B**). We then compared the Δprediction accuracy between ASD and control groups with controlling for age, sex, and site using a general linear model implemented in SurfStat (Worsley et al., 2009), and multiple comparisons across brain regions were corrected using FDR (Benjamini and Hochberg, 1995).

### Topological and network organization of prediction accuracy difference across diffusion times

We assessed underlying connectome profiles of the across-diffusion time prediction accuracy difference. First, after z-normalizing performance values of ASD individuals relative to neurotypical controls, we stratified the parcel-wise Δprediction accuracy according to seven functional communities (visual, somatomotor, dorsal attention, ventral attention, limbic, frontoparietal, default mode) (Yeo et al., 2011) and four cortical hierarchical levels (idiotypic, unimodal association, heteromodal association, paralimbic) (Mesulam, 1998) (**Fig. 3A**). Next, we associated Δprediction accuracy with a functional principal gradient, representing cortical hierarchy running from low-level sensory to higher-order transmodal system (Margulies et al., 2016). We obtained the functional gradient from the BrainSpace toolbox (Vos de Wael et al., 2020), which was generated using the Human Connectome Project database (Van Essen et al., 2013). Specifically, an affinity matrix was constructed with a normalized angle kernel with the top 10% entries for each parcel, and applied diffusion map embedding (Coifman and Lafon, 2006), which is robust to noise and computationally efficient compared to other nonlinear manifold learning techniques (Tenenbaum et al., 2000; von Luxburg, 2007). It is controlled by two parameters 𝛼 and *t*, where 𝛼 controls the influence of the density of sampling points on the manifold (𝛼 = 0, maximal influence; 𝛼 = 1, no influence) and *t* scales the eigenvalues of the diffusion operator. The parameters were set as 𝛼 = 0.5 and *t* = 0 to retain the global relations between data points in the embedded space, following prior applications (S.-J. Hong et al., 2019; Margulies et al., 2016; Paquola et al., 2019; Park et al., 2021d; Vos de Wael et al., 2020). In healthy adults, the gradient has previously been shown to follow established models of the primate cortical functional hierarchy and specifically differentiates sensory and motor networks from transmodal systems such as the default-mode network. We then associated the functional gradient with Δprediction accuracy of each individual within each group (**Fig. 3B**). The significance of the correlation was determined using 1,000 non-parametric spin-tests for accounting for spatial autocorrelation (Alexander-Bloch et al., 2018; Larivière et al., 2021; Markello and Misic, 2021). Between-group differences in the associations between ASD and control groups were assessed using two-sample t-tests with 1,000 permutation tests.

### Associations with behavioral phenotypes

As a final analysis, we investigated behavioral associations of the diffusion time-related structure-function coupling (**Fig. 4** and **Supplementary Fig. 1**). We performed multivariate analysis using PLS (Krishnan et al., 2011; McIntosh and Mišić, 2013) to associate Δprediction accuracy across diffusion times with ADOS social cognition, communication, repetitive behavior scores (Lord et al., 2000) as well as verbal and performance IQ and their ratio (verbal/performance IQ) (Hong et al., 2022). PLS is an unsupervised multivariate statistical technique that decomposes two datasets into orthogonal sets of latent variables with maximum covariance (Krishnan et al., 2011; McIntosh and Mišić, 2013). We performed PLS analysis with 1,000 bootstraps by randomly selecting subjects, and estimated PLS scores as well as loadings of the latent variables. We calculated Pearson’s correlation between the PLS scores of Δprediction accuracy and behavioral phenotypes to assess the strength of their associations. The contribution of the features which brain regions and/or behavioral phenotypes was quantified using PLS loadings. Specifically, we calculated a bootstrap ratio by dividing the mean loadings by standard errors (Zeighami et al., 2019). We thresholded the bootstrap ratio with 95% confidence interval (Zeighami et al., 2019).

### Sensitivity analysis

*a) Parcellation schemes.* We performed a structure-function coupling analysis using the Schaefer atlas with 200 parcels (Schaefer et al., 2018). To assess robustness across different spatial scales, we predicted functional connectivity from structural connectivity with the brain regions of 100 and 300 parcels (**Supplementary Figs. 2-3**).
*b) Only male participants.* Our dataset contains a larger proportion of male participants compared to female subjects. We performed the same analyses using only male participants (**Supplementary Fig. 4**).
*c) Site effects.* We obtained the data from two different sites. To assess the consistency of the results across different sites, we performed the structure-function coupling analysis for each site (**Supplementary Fig. 5**).

## Data Availability

The imaging and phenotypic data were provided, in part, by the Autism Brain Imaging Data Exchange initiative (ABIDE-II; https://fcon_1000.projects.nitrc.org/indi/abide/).

## Code Availability

The codes for simulating functional connectivity from structural connectivity are available at https://github.com/MICA-MNI/micaopen/tree/master/sf_prediction; codes for gradient generation are available at https://github.com/MICA-MNI/BrainSpace; codes for graph measures calculation are available at https://sites.google.com/site/bctnet/.

## Supplementary Information

**Table S1.** Demographic information of the study participants. Means and SDs are reported.

| Information |  | NYU |  | TCD |  | P-value |
| --- | --- | --- | --- | --- | --- | --- |
| Number<br>(Autism/Control) |  | 29/18 |  | 18/19 |  | 0.2316* |
| Age | Autism | 9.61 (6.16) | p = 0.8074 | 14.46 (3.30) | p = 0.2088 | 0.0037 |
|  | Control | 10.01 (3.95) |  | 15.83 (3.21) |  | <0.001 |
| Sex<br>(male:female) | Autism | 24:5 | p = 0.2432* | 18:0 | p = 1* | 0.0624* |
|  | Control | 17:1 |  | 19:0 |  | 0.2976* |
| ADOS – Total |  | 10.00 (3.36) |  | 8.72 (2.44) |  | 0.1924 |
| ADOS – Social cognition |  | 7.50 (2.09) |  | 5.78 (2.37) |  | 0.0226 |
| ADOS – Communication |  | 2.5 (1.70) |  | 2.94 (0.87) |  | 0.3260 |
| ADOS – Repeated<br>behavior/interest |  | 1.40 (1.27) |  | 0.22 (0.55) |  | <0.001 |
| Verbal IQ | Autism | 102.55<br>(17.07) | p = 0.0018 | 111.44<br>(13.48) | p = 0.0705 | 0.0674 |
|  | Control | 118.22<br>(13.14) |  | 119.79<br>(13.72) |  | 0.7251 |
| Performance<br>IQ | Autism | 103.28<br>(21.02) | p = 0.1399 | 109.33<br>(14.20) | p = 0.0610 | 0.2870 |
|  | Control | 112.11<br>(17.00) |  | 117.16<br>(10.16) |  | 0.2775 |
| IQ ratio | Autism | 1.01 (0.15) | p = 0.1980 | 1.02 (0.13) | p = 0.9955 | 0.6871 |
|  | Control | 1.07 (0.16) |  | 1.03 (0.13) |  | 0.3818 |

\*Chi-squared
*Abbreviations:* SD, standard deviation; NYU, New York University Langone Medical Center; TCD, Trinity College Dublin; ADOS, Autism Diagnostic Observation Schedule; IQ, intelligence quotient.

**Supplementary Fig. S1.**
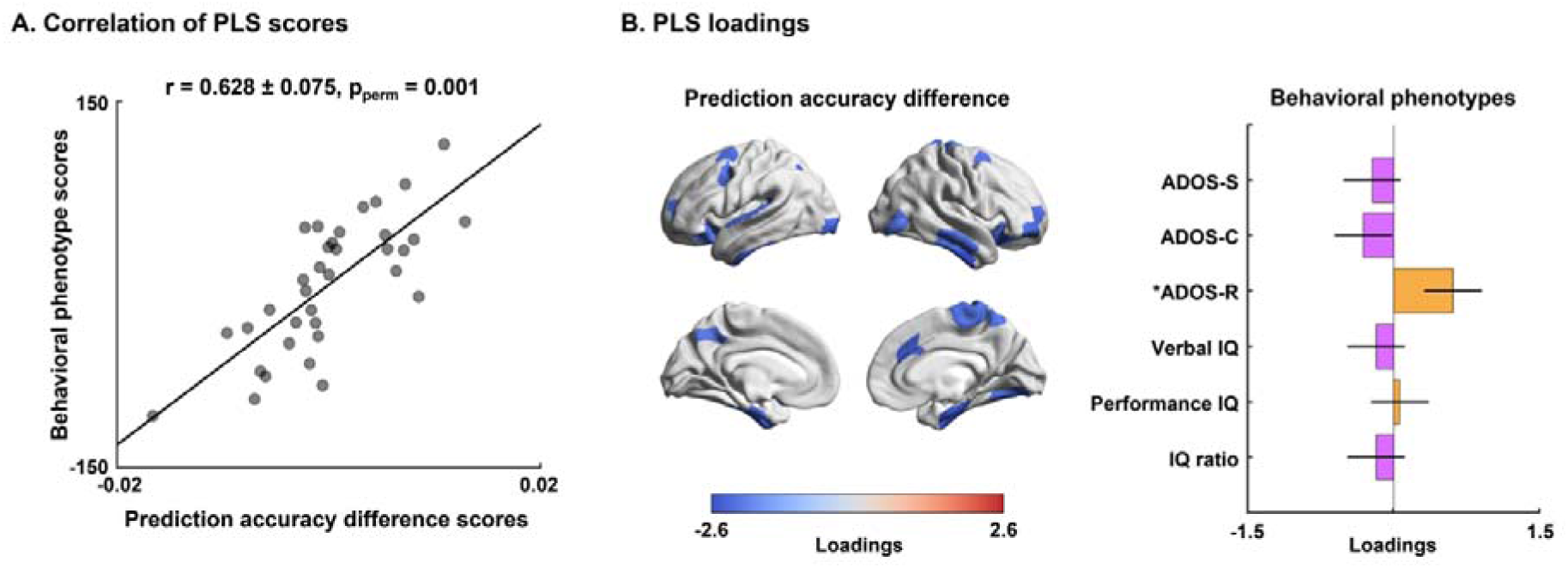
Multivariate associations between structure-function coupling imbalances and behavioral phenotypes of ASD using the second latent variable. (A) We calculated linear correlations between PLS scores of Δprediction accuracy and behavioral phenotypes of the first latent variable in ASD, which explained almost 13.8% of the variance. (B) Shown are PLS loading-based bootstrap ratios of Δprediction accuracy *(left)* and behavioral phenotypes *(right)*. Brain regions and behavioral phenotypes that showed significance are shown and marked with asterisks. *Abbreviations*: PLS, partial least squares; ADOS-S, Autism Diagnostic Observation Schedule – social cognition; ADOS-C, Autism Diagnostic Observation Schedule – communication; ADOS-R, Autism Diagnostic Observation Schedule – repeated behavior; IQ, intelligence quotient.

**Supplementary Fig. 2.**
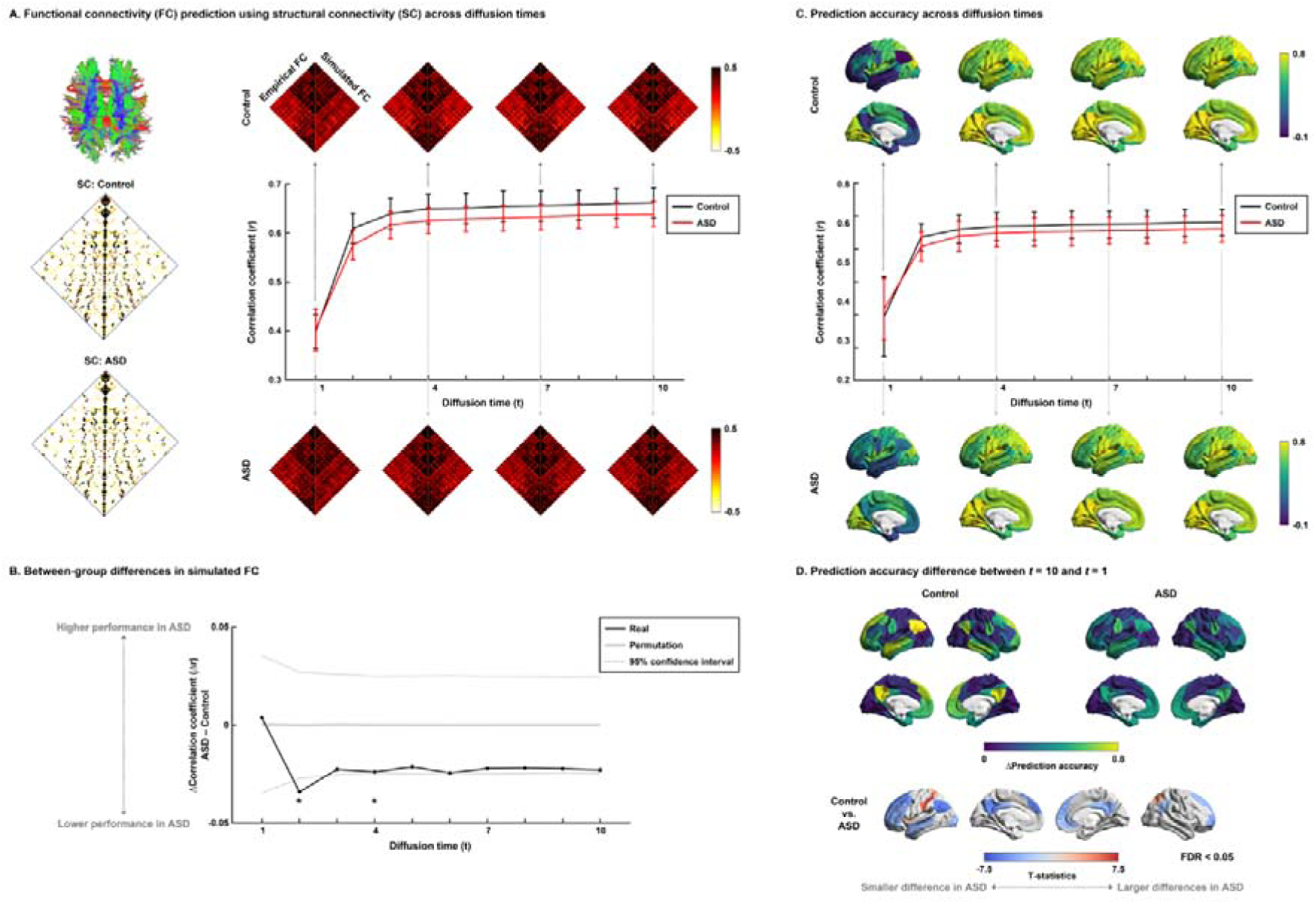
Functional connectivity prediction using structural connectivity across different diffusion times using Schaefer atlas with 100 parcels. **(A)** Group-level structural connectivity (SC) matrices in controls and ASD (*left*). Correlation coefficients between empirical and simulated functional connectivity (FC) in controls (*dark gray*) and ASD (*light gray*) as a function of diffusion time *t* (*right*). Error bars represent the SD across individuals. Shown are empirical (*left*) and simulated FC (*right*) matrices across four representative diffusion times (*t* = 1, 4, 7, and 10). **(B)** Between-group differences in prediction performance between controls and ASD. A black line indicates real differences in prediction performance between groups, a solid gray line indicates mean prediction accuracy differences across 1,000 permutation tests, and dotted gray lines indicate the 95% confidence interval. Significant between-group differences are reported with asterisks. **(C)** Correlation coefficients between empirical and simulated FC across different diffusion times *t* are shown on brain surfaces for control and ASD groups. The plot indicates correlation coefficients between empirical and simulated FC in controls (*dark gray*) and ASD (*light gray*) as a function of diffusion time. Error bars represent the SD across brain regions. **(D)** Shown are differences in prediction accuracy between the highest (*t* = 10) and lowest (*t* = 1) diffusion times (Δprediction accuracy) for both groups (*upper panels*). We assessed between-group differences in Δprediction accuracy between controls and ASDs (*lower panel*). *Abbreviation*: ASD, autism spectrum disorder; SD, standard deviation; FDR, false discovery rate.

**Supplementary Fig. 3.**
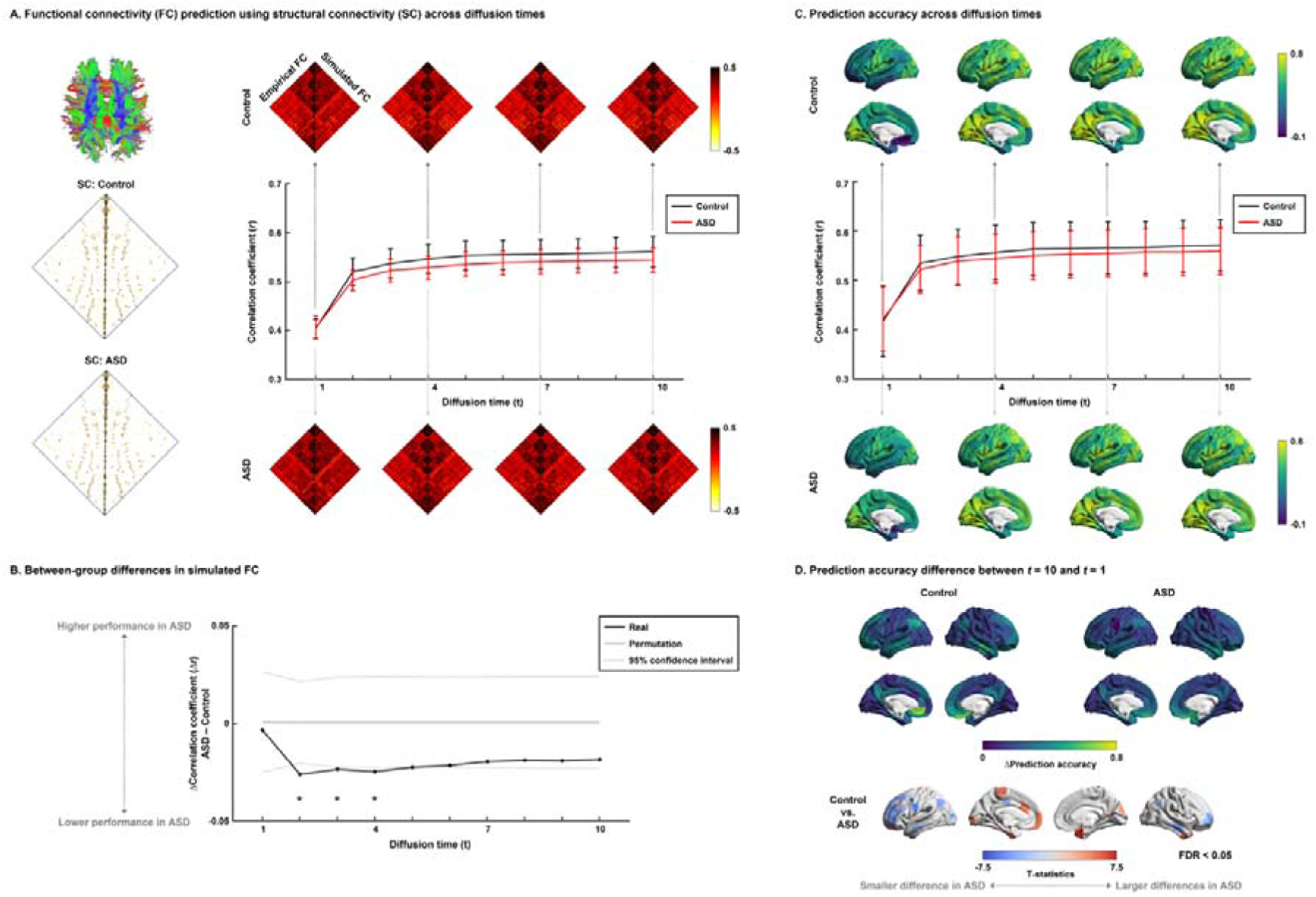
Functional connectivity prediction using structural connectivity across different diffusion times using Schaefer atlas with 300 parcels. For details, see *Supplementary Fig. 2*.

**Supplementary Fig. 4.**
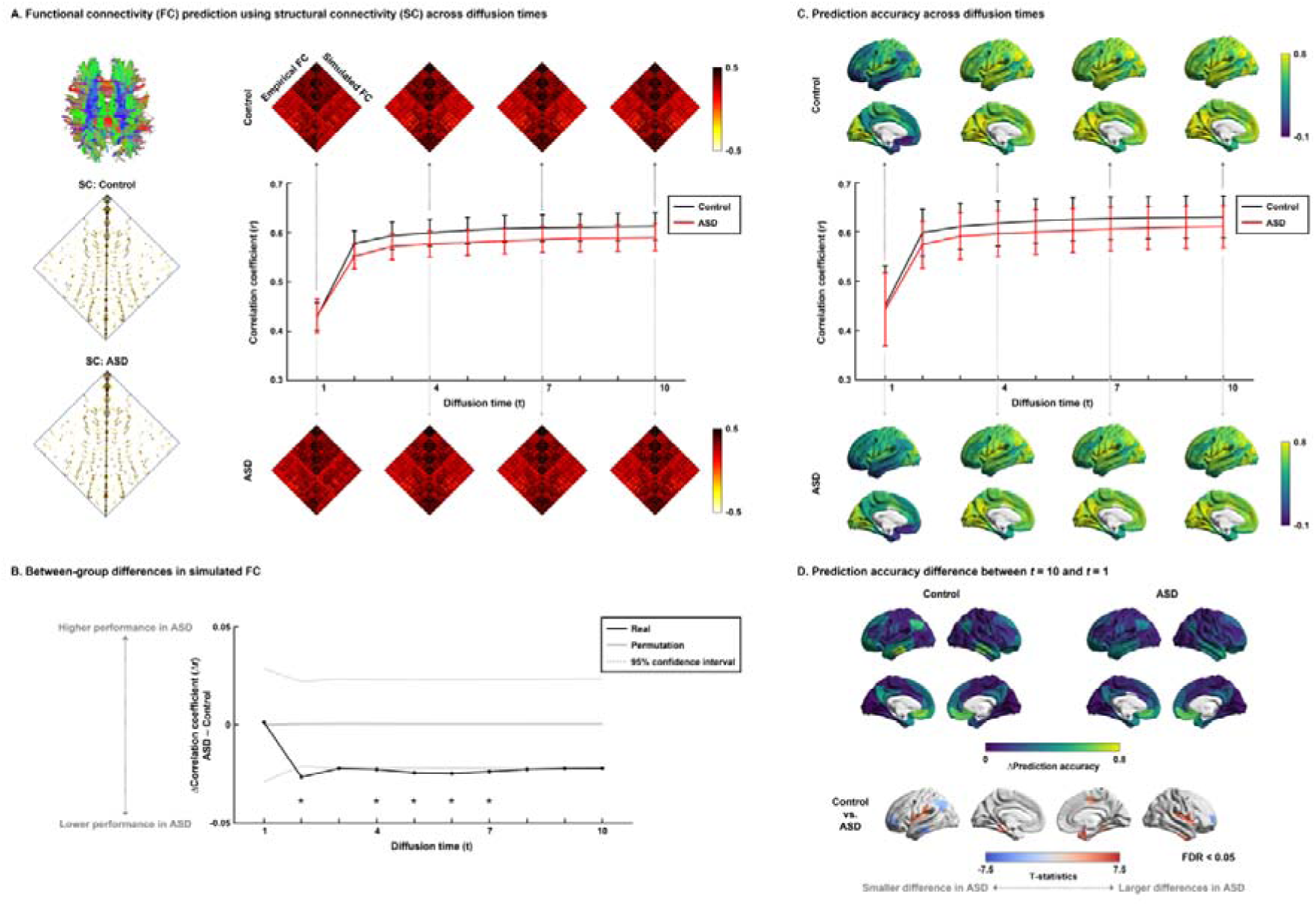
Functional connectivity prediction using structural connectivity across different diffusion times using only male participants. For details, see *Supplementary Fig. 2*.

**Supplementary Fig. S5.**
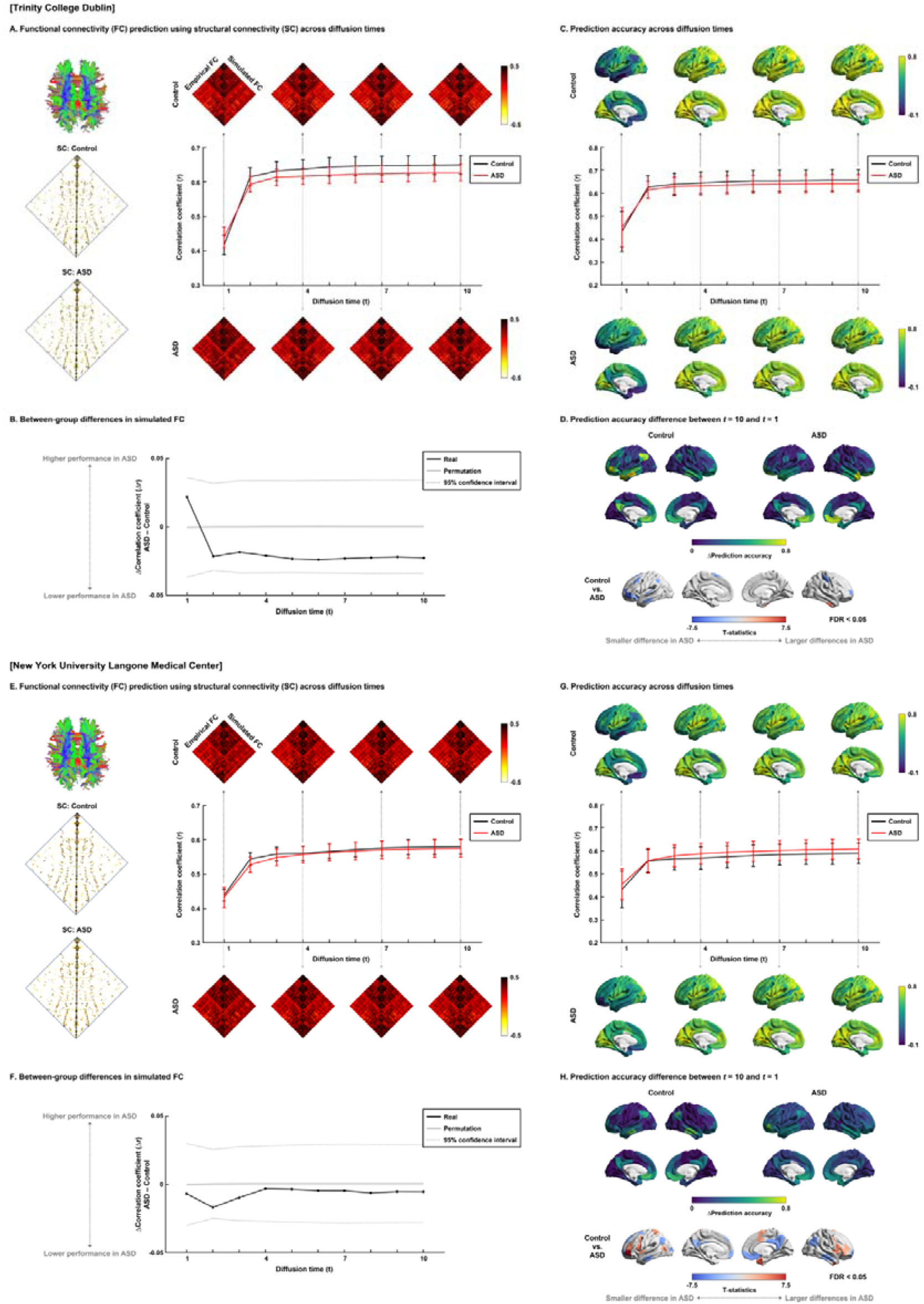
Functional connectivity prediction using structural connectivity across different diffusion times for each site separately. **(A)∼(D)** show the results from Trinity College Dublin and **(E)∼(H)** from New York University Langone Medical Center. For details, see *Supplementary Fig. 2*.

